# Exploring how the fast-slow pace of life continuum and reproductive strategies structure microorganism life history variation

**DOI:** 10.1101/2022.11.28.517963

**Authors:** Josje Romeijn, Isabel M. Smallegange

## Abstract

Studying life history strategies in microorganisms can help predict their performance when complex microbial communities can be categorised into groups of organisms with similar strategies. Microorganisms are typically classified as copiotroph or oligotroph, but it has been proven difficult to generalise their life history strategies to broad lineages. Here we tested if the fast-slow continuum and reproductive strategy framework of macro-organismal life histories can be applied to microorganisms. We used demographic and energy budget data from 13 microorganisms (bacteria, fungi, a protist and a plant) to examine how generation time, survivorship, growth form, age at maturity, recruitment success, and net reproductive rate structure microbial life histories. We found that 79% of microorganism life-history variation fell along two uncorrelated axes. Like macro-organisms, we found a fast–slow pace of life continuum, including shorter-lived microorganisms at one end, and longer-lived microorganisms that mature later in life at the other. Also, like macro-organisms, we found a second, reproductive strategy axis, with microorganisms with greater lifetime reproductive success and decreased mortality at older age at one end, and microorganisms with the opposite characteristics at the other end. Microorganismal life history strategies did not covary proportionally to their shared evolutionary history. Thus, whereas this work suggests that the macro-organismal fast-slow continuum and reproductive strategy framework could be realistically applied to microorganisms, their life history processes cannot be inferred from patterns in taxonomic composition.

**Impact statement:** Animals and plants show distinct differences in their pace of life: some have high reproduction and high mortality, others low. Here we show that microorganisms display similar such life history patterns, igniting future research on microbial life history strategies.

**Data summary:** Supplemental data, R code and MatLab code are deposited in Figshare at https://doi.org/10.6084/m9.figshare.16831543.v2 [27].

## Introduction

Different life history strategies capture different combinations of survival, growth and reproduction (demographic) rates of individuals [1] that determine the dynamics of populations. Studying life history strategies in microorganisms can help predict their performance when complex microbial communities can be categorised into organismal groups with similar strategies. Microorganisms are typically classified along a copiotroph-oligotroph dichotomy [2], where copiotrophs grow faster and rely on resource availability, but oligotrophs efficiently exploit resources at the expense of growth rate. In soils, for example, some bacterial phyla are present in greater relative abundance in response to sucrose addition, indicating copiotrophic strategies, whereas other bacterial phyla were either unresponsive or responded negatively, suggesting oligotrophic strategies [3]. Ramin and Allison [4] added a third, maintenance type to the dichotomy. Classifying microorganisms as copiotroph-oligotroph at the phylum level remains prevalent but more ecologically realistic frameworks are needed as bacterial lineages within phyla can have distinct metabolic and ecological roles [5].

In macro-organisms, the majority of life history variation is structured along (i) a fast-slow life history continuum with fast-growing, short-lived species at one end, and slow-growing, long-lived species at the other [6-8], and (ii) a reproductive strategy axis that for higher plants has iteroparous species that reproduce multiple times as adults at one end and semelparous plants that reproduce only once at the other [7,9], whereas for animals this axis is defined by the distribution of age-specific mortality hazards and the spread of reproduction [10]. We do not know if microorganismal life history variation can be structured along these same life history strategy axes [11] and if this would provide a more ecologically realistic life history framework for microbial ecologists. Such a framework could be useful because life history dynamics of individual microbial species influence processes such as human disease progression [12], human-microbe interactions [13] and even ecological strategies of coral species [14].

The aim of this explorative study is to investigate main axes of variation in microorganism life histories to offer an alternative life history framework when studying the performance of microorganisms. To this end, we first adapted a dynamic energy budget integral projection model (DEB-IPM) [15] to the microorganism life cycle. Specifically, the microorganismal DEB-IPM is a discrete-time, structured demographic model that takes data on individual cell life histories as input parameters to describe how individual cells grow according to a von Bertalanffy growth curve [16], and reproduce by binary fission to produce two offspring [17], after which individuals die. We assume that individuals have a constant background mortality. We parameterised DEB-IPMs for 13 microorganisms, including bacteria, fungi, a plant and a protist to calculate seven life history traits that inform on schedules of survival, growth, and reproduction [1]: generation time, survivorship curve, age at maturity, progressive growth (indicating size increase), retrogressive growth (indicating shrinking), mean recruitment success and net reproductive rate. We then evaluated the variation in these traits along major (life history strategy) axes using a phylogenetically corrected principal component analysis (PCA). By measuring if species life history strategies covary proportionally to their shared evolutionary history, we can assess if the fast-slow reproductive strategy framework can be used to support inferences about life history processes from patterns in taxonomic composition of microorganisms (as has been common practice by microbial ecologists [5].

## Methods

### Brief description of the DEB-IPM

The demographic function that describes growth in the microorganismal DEB-IPM [15] is derived from the Kooijman-Metz model [16], which assumes that individual (micro)organisms are isomorphic (body surface area and volume are proportional to squared and cubed length, respectively). The rate at which an individual (cell) ingests food, *I*, is assumed to be proportional to the maximum ingestion rate *I*_*max*_, the current feeding level *Y* and body surface area, and hence to the squared length of an organism: *I* = *I*_*max*_*YL*^2^. Ingested food is assimilated with a constant efficiency *ε*. A constant fraction *κ* of assimilated energy is allocated to respiration (the processes by which a cell obtains the oxygen it needs to produce energy and eliminate waste); this respiration energy equals *κεI_max_YL*^2^ and is used to first cover maintenance costs, which are proportional to body volume following *ξL*^3^(*ξ* is the proportionality constant relating maintenance energy requirements to cubed length), while the remainder is allocated to somatic growth. This means that, if an individual cell survives from hour *t* to hour *t* + 1, it grows from length *L* to length *L*’ following a von Bertalanffy growth curve, 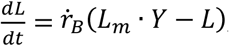, where 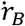 is the von Bertalanffy growth rate *L_m_* = *κεI_max_*/*ξ* and is the maximum length under conditions of unlimited resource. Both *κ* and *I*_*max*_ are assumed to be constant. However, the role of *κ* in the DEB-IPM is mostly implicit, as *κ* is used as input parameter only in the starvation condition (see below), whereas *L*_*m*_ is measured directly from data.

Growth, as described above, as well as survival and reproduction are captured in the DEB-IPM by four fundamental functions to describe the dynamics of a well-mixed population comprising cohorts of individual cells of different sizes [15]: (1) the survival function, *S*(*L*(*t*)) (unit: h^−1^), describing the probability of surviving from hour *t* to hour *t*+1; (2) the growth function, *G*(*L*′,*L*(*t*)) (unit: h^−1^), describing the probability that an individual of body length *L* at hour *t* grows to length *L’* at *t* + 1, conditional on survival; (3) the reproduction function, *R*(*L*(*t*)) (unit: # h^−1^), giving the number of offspring produced between hour *t* and *t* + 1 by an individual of length *L* at hour *t*; and (4) the parent-offspring function, *D*(*L*′,*L*(*t*)) (unit: h^−1^), the latter which describes the association between the body length of the parent cell *L* and offspring cell length *L*’ (i.e. to what extent does offspring size depend on parental size). Denoting the number of cells at hour *t* by *N*(*L,t*) means that the dynamics of the cell length number distribution from hour *t* to *t* + 1 can be written as:

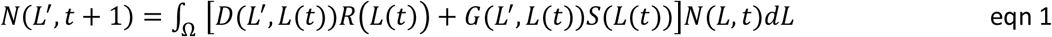

where the closed interval Ω denotes the cell length domain. Note that this simple DEB-IPM does not account for temperature effects or reserve dynamics (which can, however, be an acceptable simplification [18]). Implicitly underlying the population-level model of eqn 1, like in any IPM, is a stochastic, individual-based model, in which individuals follow Markovian growth trajectories that depend on an individual’s current state (Easterling et al. 2000). This individual variability is in standard IPMs modelled in the functions describing growth, *G*(*L*′,*L*(*t*)), and the parent-offspring association, *D*(*L*′,*L*(*t*)), using a probability density distribution, typically Gaussian [19]. In the DEB-IPM, this individual variability arises from how individuals experience the environment; specifically, the experienced feeding level *Y* follows a Gaussian distribution with mean *E*(*Y*) and standard deviation *σ* (*Y*). It means that individual cells within a cohort of length *L* do not necessarily experience the same feeding level due to demographic stochasticity (e.g. individuals, independently of each other, have good or bad luck in their feeding experience).

The survival function *S*(*L*(*t*)) in eqn 1 is the probability that an individual of cell length *L* survives from hour *t* to *t* + 1:

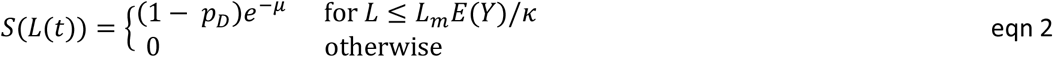

assuming a constant background mortality rate *μ* (h^−1^). The parameter *L*_*m*_ (μm) is the maximum cell length under conditions of unlimited abundance of food of the highest, relative quality, *E*(*Y*) = 1. The probability of an individual dividing, and subsequently produce *n*_*O*_ = 2 offspring, *p*_*D*_, enters the survival function to represent binary fission [17], because reproduction in microorganisms entails that the dividing individual ceases to exist. Individuals die from starvation at a cell length at which maintenance requirements exceed the total amount of assimilated energy, which occurs when *L* > *L_m_*·*E*(*Y*)/*κ* and hence, then, *S*(*L*(*t*)) =0 (e.g., an individual of size *L*_*m*_ will die of starvation if *E*(*Y*) < *κ*).

The function *G*(*L*′,*L*(*t*)) is the probability that an individual of cell length *L* (μm) at hour *t* grows to length *L’* at *t* + 1, conditional on survival, and follows a Gaussian distribution [19]:

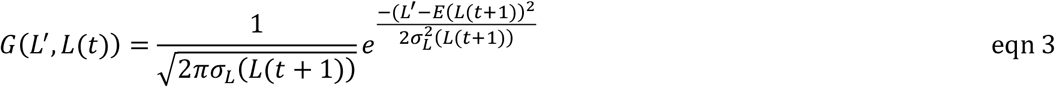

With

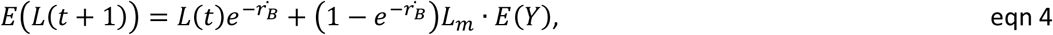

And

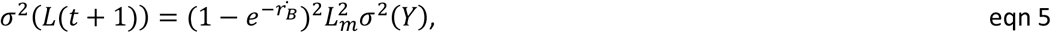

where 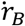 is the von Bertalanffy growth rate and *σ*(*Y*) is the standard deviation of the expected feeding level, assuming individuals can shrink under starvation conditions.

The reproduction function *R*(*L*(*t*)) follows a ‘pre-breeding’ census as cell division, i.e. reproduction, is fatal. *R*(*L*(*t*)) gives the number of offspring produced between hour *t* and *t* + 1 by an individual of cell length *L* (μm) at hour *t*:

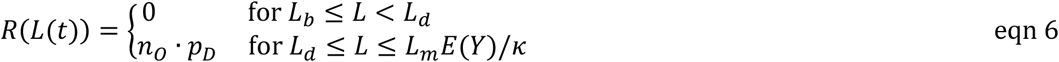

An individual always divides to produce *n*_*O*_ = 2 offspring [one parent and one offspring cell]. We set the cell length at which individuals divide, *L*_*d*_ (μm), equal to the asymptote of the von Bertalanffy growth curve in cell length so that *L_d_* = *E*(*Y*)·*L_m_*. Only surviving adults reproduce; thus, only individuals within a cohort of cell length *L_d_* ≤ *L* ≤ *L_m_Y*/*κ* reproduce. The probability of an individual dividing, *p*_*D*_, equals:

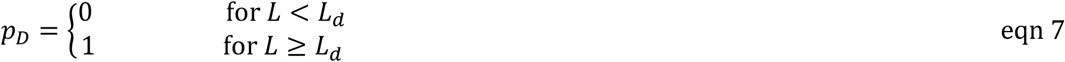

Note that the probability of dividing also enters the survival function as division is fatal.

The probability density function *G*(*L*′,*L*(*t*)) gives the probability that the offspring of an individual of cell length *L* are of length *L′* at hour *t* + 1:

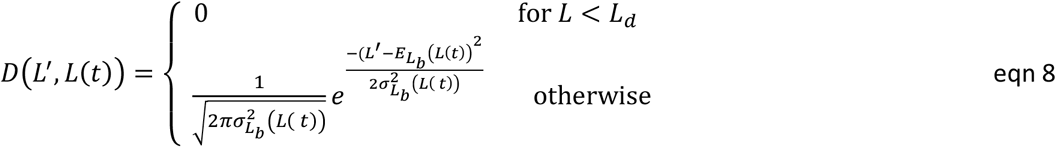

Where 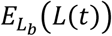 is the expected cell length of offspring produced by a cohort of individuals with length *L*(*t*), and 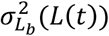 the associated variance. For simplicity, we set 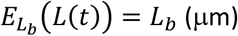 constant, and assumed its associated variance, 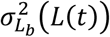, to be negligibly small so that offspring within a species are evenly sized.

### Parameterization of the DEB-IPM

The projection interval between time *t* and time *t* + 1 was set at 1 hour; all rates and durations are therefore expressed per hour, or in hours, respectively. We parameterized DEB-IPMs for 13 microorganisms (Fig. 1) (Table 1), with constant values for κ = 0.9 (for macro-organisms, κ is assumed to be 0.8; we assumed this would be higher for microorganisms as most energy goes into respiration and not reproduction), μ = 0.01 h^−1^ (assuming background mortality under laboratory circumstances is very low), and *n*_O_ = 2 (as one microorganism divides to produce two offspring) (Table 1). Values of *L*_*b*_, *L*_*d*_, *L*_*m*_, and 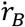 were taken from the literature (references in Table 1) or measures of cell length at birth and the elongated length of a single cell were combined to give *L*_*m*_. In case of *Pseudomonas aeruginosa*, only log values of cell length were available [22], which we backtransformed. Cell length measures of *Bacillus subtilis* were available for different media, and we used those values obtained for cells grown on a favourable, glycerol rich medium [23]. In case of *Dictyostelium discoideum*, we inferred from photos that cells are circle-shaped in between birth and mitosis [24], so that cell surface *S* = *πr*^2^, from which follows that the cell length equals 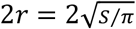. Cell surfaces measures to calculate cell lengths of *D. discoideum* were taken from two timeframe photos taken between mitosis and birth [24] Finally, we manually measured cell lengths of *Desulfovibrio vulgaris* from a series of scale-barred images of cell length over time [25]. For all species, we estimated the von Bertalanffy growth rate, 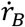, using a nonlinear least-squares estimation procedure in R (nls function) [26] to fit the equation 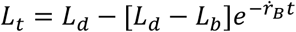 for 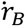 [15] against observed growth curves of cell length *L*_*t*_ (μm) over time *t* (hours), taken from the literature (see R code in [27]).

**Table 1.** Model parameters for the 13 microorganisms (Fig. 1). Parameters are expected feeding level *E*(*Y*), the fraction of assimilated energy allocated to respiration, κ, cell length at birth, *L*_*b*_, length at division, *L*_*d*_, maximum length, *L*_*m*_, mortality rate, μ, von Bertalanffy growth rate,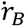, and number of offspring produced, *n*_*O*_. Units are in brackets. Superscript numbers are references. Species numbers are the same as in the Fig. 1. See further main text.

| ID | Species (kingdom) | Model parameters |  |  |  |  |  |  |  |
| --- | --- | --- | --- | --- | --- | --- | --- | --- | --- |
| | | $E(Y)$<br>(–) | $\kappa$<br>(–) | $L_b$<br>($\mu\text{m}$) | $L_d$<br>($\mu\text{m}$) | $L_m$<br>($\mu\text{m}$) | $\mu$<br>($\text{h}^{-1}$) | $n_o$<br>(#) | $\dot{r}_B$<br>($\text{h}^{-1}$) |
| 1 | *Dictyostelium discoideum*<br>(Protista) | 0.80 | 0.90 | 15.26<br>[24] | 24.74<br>[24] | 48.48<br>[24] | 0.01 | 2 | 0.12 |
| 2 | *Chlamydomonas reinhardtii*<br>(Plants) | 1.00 | 0.90 | 12.64<br>[73] | 27.26<br>[73] | 29.23<br>[73] | 0.01 | 2 | 0.03 |
| 3 | *Bacillus subtilis*<br>(Bacteria) | 0.80 | 0.90 | 3.84<br>[23] | 6.89<br>[23] | 13.87<br>[23] | 0.01 | 2 | 1.62 |
| 4 | *Synechococcus elongatus*<br>(Bacteria) | 1.00 | 0.90 | 2.04<br>[72] | 5.98<br>[72] | 8.031<br>[72] | 0.01 | 2 | 0.03 |
| 5 | *Mycobacterium smegmatis*<br>(Bacteria) | 0.80 | 0.90 | 5.06<br>[64] | 8.72<br>[64] | 11.96<br>[65] | 0.01 | 2 | 0.09 |
| 6 | *Pseudomonas aeruginosa*<br>(Bacteria) | 0.80 | 0.90 | 3.86<br>[22] | 17.34<br>[22] | 26.00<br>[68] | 0.01 | 2 | 0.25 |
| 7 | *Shewanella oneidensis*<br>(Bacteria) | 0.80 | 0.90 | 1.35<br>[69] | 3.35<br>[69] | 5.00<br>[69] | 0.01 | 2 | 0.17 |
| 8 | *Escherichia coli* (Bacteria) | 0.50 | 0.90 | 2.18<br>[66] | 3.46<br>[66] | 8.00<br>[67] | 0.01 | 2 | 5.21 |
| 9 | *Caulobacter vibrioides*<br>(Bacteria) | 0.70 | 0.90 | 1.76<br>[70] | 3.22<br>[70] | 5.30<br>[71] | 0.01 | 2 | 1.10 |
| 10 | *Methylobacterium extorquens*<br>(Bacteria) | 0.90 | 0.90 | 1.71<br>[74] | 4.40<br>[74] | 5.24<br>[74] | 0.01 | 2 | 0.15 |
| 11 | *Desulfovibrio vulgaris*<br>(Bacteria) | 1.00 | 0.90 | 3.15<br>[25] | 6.18<br>[25] | 6.19<br>[25] | 0.01 | 2 | 0.49 |
| 12 | *Schizosaccharomyces pombe*<br>(Fungi) | 0.90 | 0.90 | 8.67<br>[61,79] | 16.00<br>[61,79] | 19.43<br>[62] | 0.01 | 2 | 0.69 |
| 13 | *Saccharomyces cerevisiae*<br>(Fungi) | 0.99 | 0.90 | 3.78<br>[37] | 8.12<br>[37] | 8.24<br>[20] | 0.01 | 2 | 0.56 |

estimated from other data. Specifically, in case of *S. cerevisiae*, we calculated cell length as the cell diameter $2r$ ( $\mu\text{m}$ ) from cell volume $V$ ( $\mu\text{m}^3$ ), assuming *S. cerevisiae* cells are sphere-shaped so that $V = \frac{4}{3}\pi r^3$ , from which follows that cell length equals $2r = 2\left(\frac{3}{4\pi}V\right)^{1/3}$ . To calculate $L_m$ for *S. cerevisiae*, we used the volume of a mother cell and bud [20], and subtracted the volume of the mother cell at division from the volume of the mother cell at birth, assuming that the size at birth of the mother equals the bud's size. Cell length of the mother and bud were calculated from the volume measures as before, and added together to give maximum length, $L_m$ , of *S. cerevisiae*. In case of *Chlamydomonas reinhardtii*, we assumed that, given their shape, cells were proportioned as $4 \times 4 \times 10 \mu\text{m}$ [21]. Given the fact that cell volume $V = \frac{4}{3}\pi r^3$ and $r^3 = 4a \cdot 4a \cdot 10a$ , gives cell length as $10a = \sqrt[3]{3V/(640\pi)}$ , where $a$ is the ratio between the 3 dimensions of the cell, making '10a' the longest cell dimension and therefore interpretable as cell length. In case of *Mycobacterium smegmatis*,

**Fig. 1.**
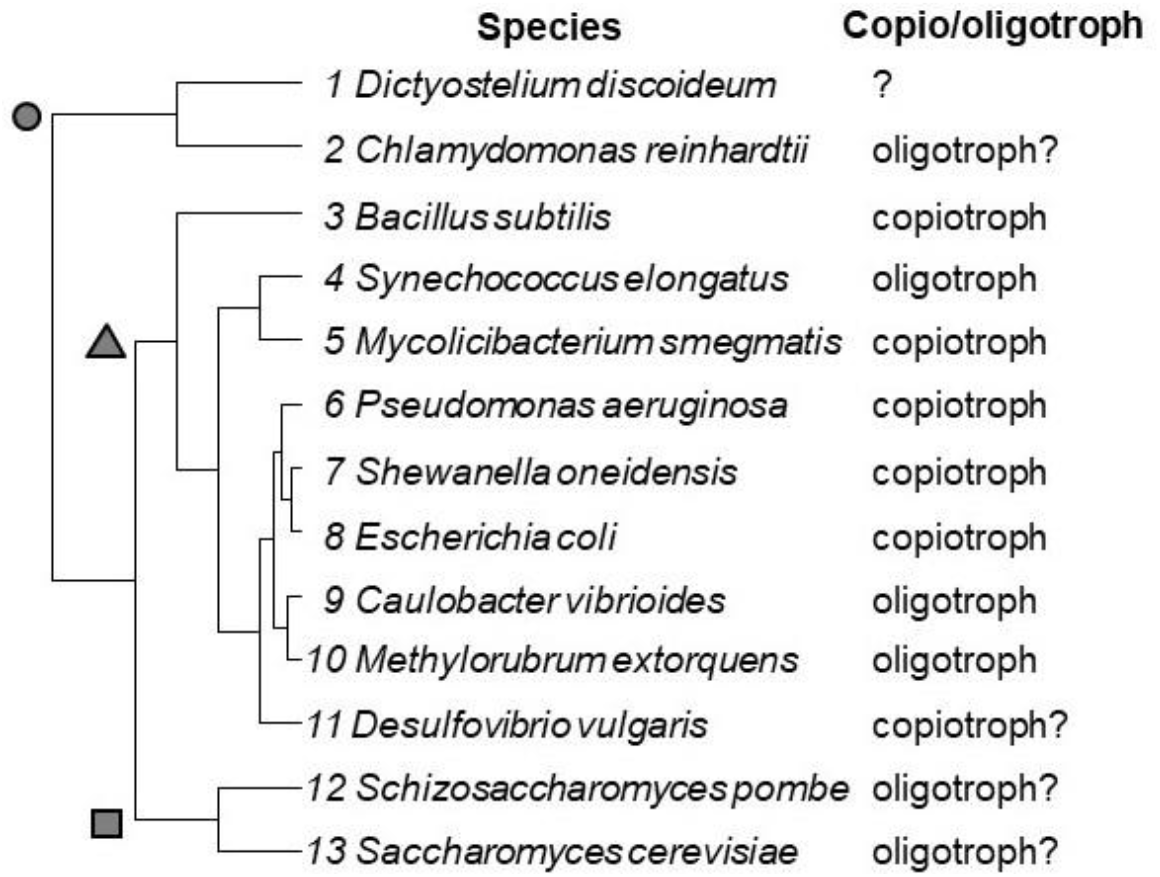
Phylogenetic tree of the 13 study species created using the *rotl* package [54] to download a tree from the Open Tree of Life (OTL) [55], after taxonomical updating using the R library *taxize* [56] (input names of *Synechococcus elongatus, P. aeruginosa* and *Escherichia coli* were changed to *Synechoccocus elongatus* CCMP1630, *Pseudomonas aeruginosa* 148 and *Escherichia coli* 402290, respectively). Branch length was computed using the *compute*.*brlen* in the R package *ape* [57]. Polytomies were collapsed using the function *multi2di* in *ape* [55]. Copiotroph and oligotroph classification was according to Stone et al. 2023 for species 2-10, and suspected classifications (indicated with questions marks) are from [58] (species 2), [59] (species 11), and [60] (species 12-13).

### Identifying life history patterns

For each species, we used their DEB-IPM to calculate in MATLAB [version 9.10.01602886 R2021a] seven representative life history traits: generation time *T*, the shape of the survivorship curve *H*, age at maturity *L*_α_, progressive growth γ, retrogressive growth ρ, mean recruitment success φ, and net reproductive rate R_0_ [9] (Table 2) that inform on schedules of survival, growth and reproduction. Because all individuals die after reproducing, we did not include mature life expectancy or degree of iteroparity [9]. We set *E*(*Y*) for each species such that it mimics favourable laboratory conditions and exponential growth, so that λ > 1 and *R*_*0*_ > 1 (Table 3). We evaluated the variation in these traits along major axes using phylogenetically corrected principal component analysis (p-PCA) [28] conducted in R [26] to identify the major life history strategy axes. To this end we used the *phyl*.*pca* function in the R library *phytools* [7,9,28], after log-transforming and scaling values to μ = 0 and SD = 1 to agree with PCA assumptions.

**Table 2.** Life history traits and their loadings onto the first two PC axes. To calculate life history traits, we discretised the IPM (Eqn 1) by dividing the length domain Ω into 200 very small-width discrete bins, resulting in a matrix **A** of size *m* × *n*, where *m* = *n* = 200, and which dominant eigenvalue equals λ. Mean lifetime reproductive success *R*_0_ is the dominant eigenvalue of the matrix **F = V(I − GS)**^−1^, where **I** is the identity matrix and **V** = **DR**, with **D** as the parent-offspring association, **R** the reproduction, **G** the growth and **S** the survival matrix (Caswell 2001); this gives generation time *T* = log(*R*_0_)/log(*λ*) [75]. The mean life expectancy, *η*_e_, is calculated as *η*_e_ = **1**^T^**N**, where **1** is a vector of ones of length *m* and **N** is the fundamental matrix **N = (I − S)**^−1^. The longevity of an individual of length *L* is *η*_L_, which means we can calculate age at sexual maturity 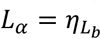 and mature life expectancy 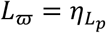 so that 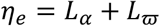 [76: eqn 4.21]. *l*_x_ is the probability of surviving to age at least *x*, and *m*_x_ is the average fertility of age class *x* (cf. [77]. 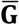 the mean of **G**,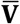 the mean of **V**, and *i* and *j* are the row and column entries of the matrix, respectively. The vital rates included in the studied set of fundamental life history traits (progressive growth *γ*, retrogressive growth *ρ*, and sexual reproduction *φ*) were averaged across the columns *j* (the length bins), weighted by the relative contributions of each stage at stationary equilibrium. For example, to calculate mean sexual reproduction *φ*, we summed the values in the columns *j* of the **V** matrix and multiplied each *φ*_*ij*_ by the corresponding *j*th element *w*_j_ of the stable stage distribution **w**, calculated as the right eigenvector of **A**.

| Life history trait | Symbol | Definition | Equation | p-PC1 | p-PC2 | p-PC3 | PC1 | PC2 | PC3 |
| --- | --- | --- | --- | --- | --- | --- | --- | --- | --- |
| Generation time | $T$ | Number of hours required for the individuals of a population to be replaced by new ones | $T = \frac{\log(R_0)}{\log(\lambda)}$ | **0.98** | -0.02 | -0.08 | 0.48 | -0.02 | -0.07 |
| Survivorship curve | $H$ | Keyfitz' entropy ($H < 1$ denotes increasing mortality rate with age, $H > 1$ denotes decreasing mortality rate with age, $H = 1$ denotes a constant mortality rate with age) [78]. | $H = -\frac{\sum_{x=0}^{x=\eta_e} \log(l_x) l_x}{\sum_{x=0}^{x=\eta_e} l_x}$ | 0.45 | **-0.78** | -0.33 | 0.22 | **-0.70** | -0.31 |
| Age at maturity | $L_\alpha$ | Number of hours for an individual in the population to become reproductive | $L_\alpha = \eta_{L_b}$ | **0.98** | -0.06 | -0.06 | 0.48 | -0.05 | -0.06 |
| Progressive growth | $\gamma$ | Mean probability of growing to a larger length across the length domain $\Omega$. | $\gamma = \sum_i^m \bar{\mathbf{G}}_{i,j} \bar{\mathbf{w}}_j \vert _{i < j}$ | **0.92** | 0.24 | 0.05 | 0.44 | 0.21 | 0.04 |
| Retrogressive growth | $\rho$ | Mean probability of growing to a smaller length across the length domain $\Omega$. | $\rho = \sum_i^m \bar{\mathbf{G}}_{i,j} \bar{\mathbf{w}}_j \vert _{i > j}$ | **-0.53** | -0.25 | **-0.78** | -0.26 | -0.22 | **-0.73** |
| Mean recruitment success | $\varphi$ | Mean per-capita number of recruits across the length domain $\Omega$. | $\varphi = \sum_i^m \bar{\mathbf{V}}_{i,j} \bar{\mathbf{w}}_j$ | **-0.99** | 0.04 | 0.07 | -0.48 | 0.04 | 0.06 |
| Net reproductive rate | $R_0$ | Mean number of recruits produced during the mean life expectancy of an individual in the population. | $R_0 = \sum_{x=0}^{x=\eta_e} l_x m_x$ | -0.16 | **-0.72** | **0.65** | -0.08 | **-0.65** | **0.60** |

Loadings in bold indicate a high contribution (greater than $\pm 0.50$ ) of the life-history trait to the PC axis. p-PC axes are from the phylogenetically constructed PCA, and PC axes from a 'normal' PCA.

**Table 3.** The life history traits generation time *T*, survivorship curve *H*, age at sexual maturity *L_α_*, progressive growth γ, retrogressive growth ρ, mean sexual reproduction *φ*, and net reproductive rate *R*_0_ calculated from DEB-IPMs and analysed using PCAs (Table 2). Species numbers are the same as in Fig. 1.

| Species | Life history trait |  |  |  |  |  |  |
| --- | --- | --- | --- | --- | --- | --- | --- |
| | $T$ | $H$ | $L_\alpha$ | $\gamma$ | $\rho$ | $\phi$ | $R_0$ |
| 1 *Dictyostelium discoideum* | 6.11 | 0.11 | 5.69 | 4.29E-05 | 3.11E-08 | 0.24 | 1.69 |
| 2 *Chlamydomonas reinhardtii* | 73.80 | 0.27 | 51.90 | 1.31E-04 | 2.35E-06 | 0.02 | 0.96 |
| 3 *Bacillus subtilis* | 2.01 | 0.02 | 2.00 | 3.21E-05 | 3.49E-06 | 0.82 | 1.16 |
| 4 *Synechococcus elongatus* | 37.54 | 0.15 | 31.00 | 1.73E-04 | 2.65E-09 | 0.04 | 1.32 |
| 5 *Mycobacterium smegmatis* | 20.65 | 0.22 | 18.35 | 4.72E-05 | 2.75E-06 | 0.07 | 1.49 |
| 6 *Pseudomonas aeruginosa* | 7.90 | 0.17 | 7.63 | 3.83E-05 | 8.32E-07 | 0.18 | 1.71 |
| 7 *Shewanella oneidensis* | 10.04 | 0.18 | 9.50 | 3.72E-05 | 9.17E-07 | 0.14 | 1.71 |
| 8 *Escherichia coli* | 2.27 | 0.16 | 2.32 | 2.13E-05 | 1.93E-05 | 0.71 | 1.20 |
| 9 *Caulobacter vibrioides* | 2.88 | 0.21 | 2.89 | 2.86E-05 | 6.99E-06 | 0.54 | 1.39 |
| 10 *Methylobacterium extorquens* | 16.45 | 0.23 | 15.20 | 4.48E-05 | 2.07E-06 | 0.09 | 1.57 |
| 11 *Desulfovibrio vulgaris* | 8.59 | 0.34 | 8.55 | 5.78E-05 | 5.40E-06 | 0.17 | 1.52 |
| 12 *Schizosaccharomyces pombe* | 4.24 | 0.22 | 4.20 | 3.87E-05 | 3.86E-06 | 0.35 | 1.50 |
| 13 *Saccharomyces cerevisiae* | 7.79 | 0.33 | 7.81 | 5.74E-05 | 5.34E-06 | 0.18 | 1.51 |

We estimated Pagel’s λ to inform on the phylogenetic correlation between species traits expected under Brownian motion [29], which varies between 0 (phylogenetic independence) and 1 (species traits covary proportionally to their shared evolutionary history). We used Pagel’s λ because it has been used successfully in previous cross-taxonomical PCAs on life history traits [7-9], it can describe the evolution of bacterial trait distributions well [30], and suffers from low type I and II errors for phylogenies of varying sizes [31]. Pagel’s λ is estimated jointly for all life history traits [28]. Note that Pagel’s λ does not detect non-Brownian models of evolution, which can yield phylogenetic dependence (λ = 1). Intermediate levels of phylogenetic dependence (0 < λ < 1) could result from the Ornstein-Uhlenbeck mode of evolution (e.g, [32,33]), in which phylogenetic influence decays through time as species trait values move toward an evolutionary optimum irrespective of the starting point [29]. In the simplest, non-Brownian model, traits may evolve instantaneously in response to selection, in which case traits would show no phylogenetic dependence (λ = 0). Additional tests are required to distinguish these forms of models. Since Pagel’s λ was close to 0 (see Results), we also conducted a normal PCA on the life history traits. We assessed the significance of PC axes using Kaiser’s criterion [34]).

## Results

The range of microorganismal life history traits was captured by the first three p-PC axes of the p-PCA, which together explained 79% of the variation (Fig. 2A). The life history traits most closely aligned with p-PC1 are related to the fast-slow continuum [1]: generation time (*T*), age at maturity (*L*_α_) and mean recruitment success (φ) had the greatest loadings onto p-PC1. The positive loading of generation time onto p-PC1 had an opposite loading for mean recruitment success, supporting a trade-off between highly reproductive species and population turnover [1] (Table 2). The rate of individual cell growth (progressive growth, γ) and shrinkage (retrogressive growth, ρ) also loaded onto p-PC1 (Table 2). Thus, as we move from negative to positive scores along p-PC1, individuals increase their allocation to individual growth and to the longevity-related life history trait age at maturity and decrease in population turnover (i.e., greater generation time) at the expense of the production of new recruits (Fig. 2A). Only at the extreme ends of this pace of life continuum was there a match between copiotrophs at the fast end and oligotrophs at the slow end of the continuum (red (copiotroph) and blue (oligotroph) convex hulls do not overlap at the extreme ends in Fig. 2A).

**Fig. 2.**
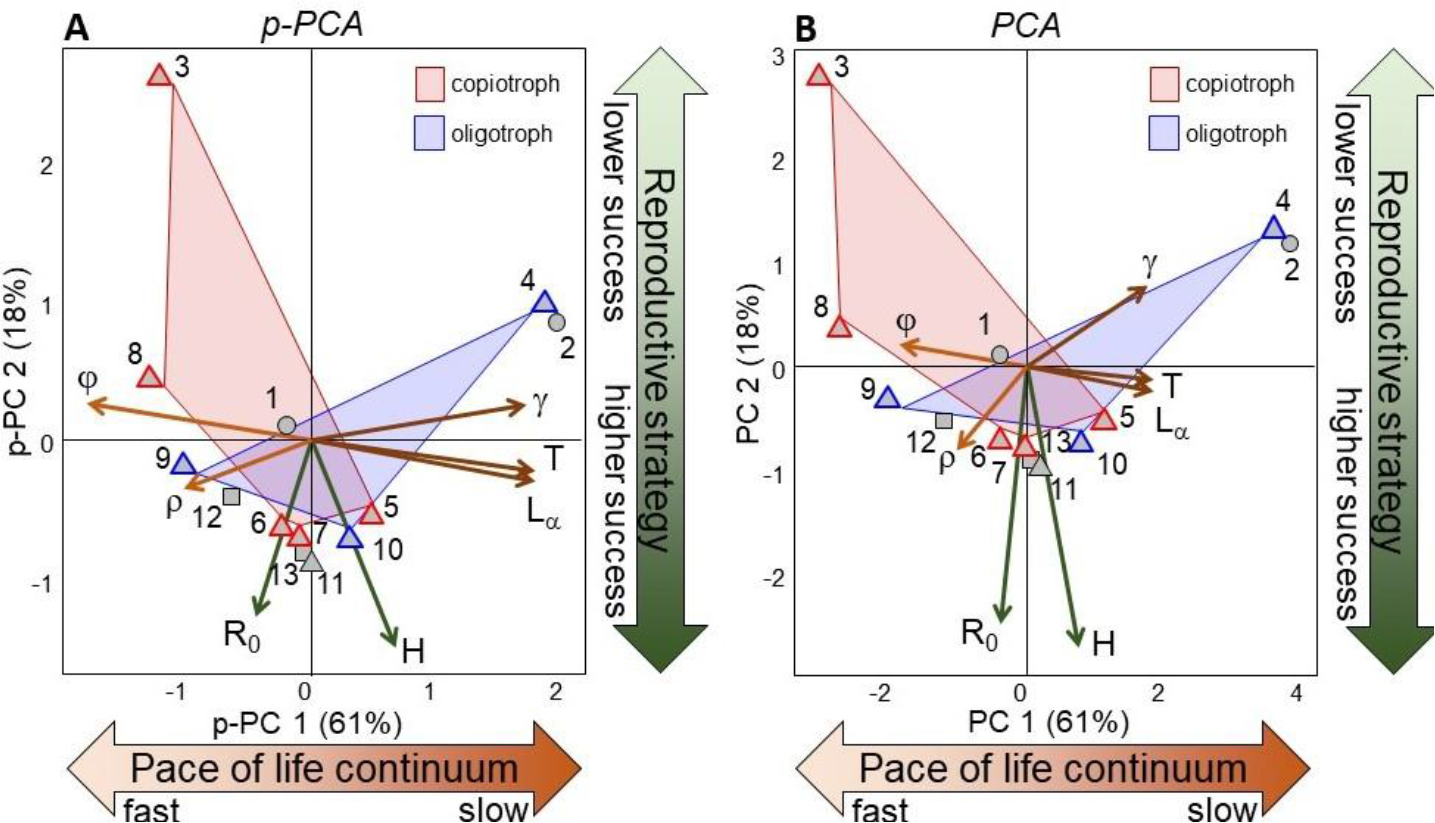
To characterise life histories, a phylogenetically corrected PCA (p-PCA) (**A**) and a normal PCA (**B**) were performed on generation time *T*, the shape of the survivorship curve *H*, age at maturity *L*_α_, progressive growth γ, retrogressive growth ρ, mean recruitment success φ, and net reproductive rate R_0_ (Table 3). Both show the same result. We show the first two PC axes, although the eigenvalue of the third PC axis was also > 1 (following the Kaiser criterion [34]: λ_axis 1_ = 4.27; λ_axis 2_ = 1.25; λ_axis 3_ = 1.15; λ_axis 4_ = 0.25) and the third axis explained 16% of the variation (see further main text). Arrow lengths are proportional to the loadings of each trait onto these two axes. Circles indicate protist or plant, triangles indicate bacteria, and squares indicate fungi. Numbers next to symbols indicate species identity as in Fig. 1 and in Tables 1 and 3. Confirmed classifications of species as copiotroph (red-lined symbols) and oligotroph (blue-lined symbols) (Fig. 1) are indicated in both biplots and the associated convex hulls denote areas of life history strategy space occupied by them.

The two life history traits aligned with p-PC2 represent dimensions of microorganism reproductive strategy not captured by mean recruitment success: both the net reproductive rate (*R*_0_) and the shape of the survivorship curve (*H*) are negatively loaded onto p-PC2: the reproductive strategy axis (Table 2). This means that, as we move from positive to negative scores along p-PC2, microorganisms attain greater lifetime reproductive success while the mortality of older individuals decreases. This time, species classified as copiotrophs and oligotrophs completely overlapped along the reproductive strategy axis (red and blue convex hulls in Fig 2A, respectively).

Interestingly, p-PC3 is retained [its associated eigenvalue >1 (Fig. 2A)] and explained 16% of the variation. The negative loading of net reproductive rate R_0_ onto p-PC3 had an opposite loading for retrogressive growth ρ (Table 2): moving from one end of p-PC3 to the other, microorganisms attain greater (or lower) lifetime reproductive success and tend to shrink less (or more).

Finally, there was a weak phylogenetic signature in the structuring of life history variation (Pagel’s λwas close to zero: λ < 0.01), suggesting a weak role of overall phylogenetic ancestry in life history trait variation, confirmed by traitgrams that showed no strong mapping of a species’ place along the pace of life continuum (Fig. 3A) or reproductive strategy axis (Fig. 3B) and its phylogenetic position. We therefore ran a normal PCA, which showed the same results as the p-PCA (Table 3, Fig. 2B).

**Fig. 3.**
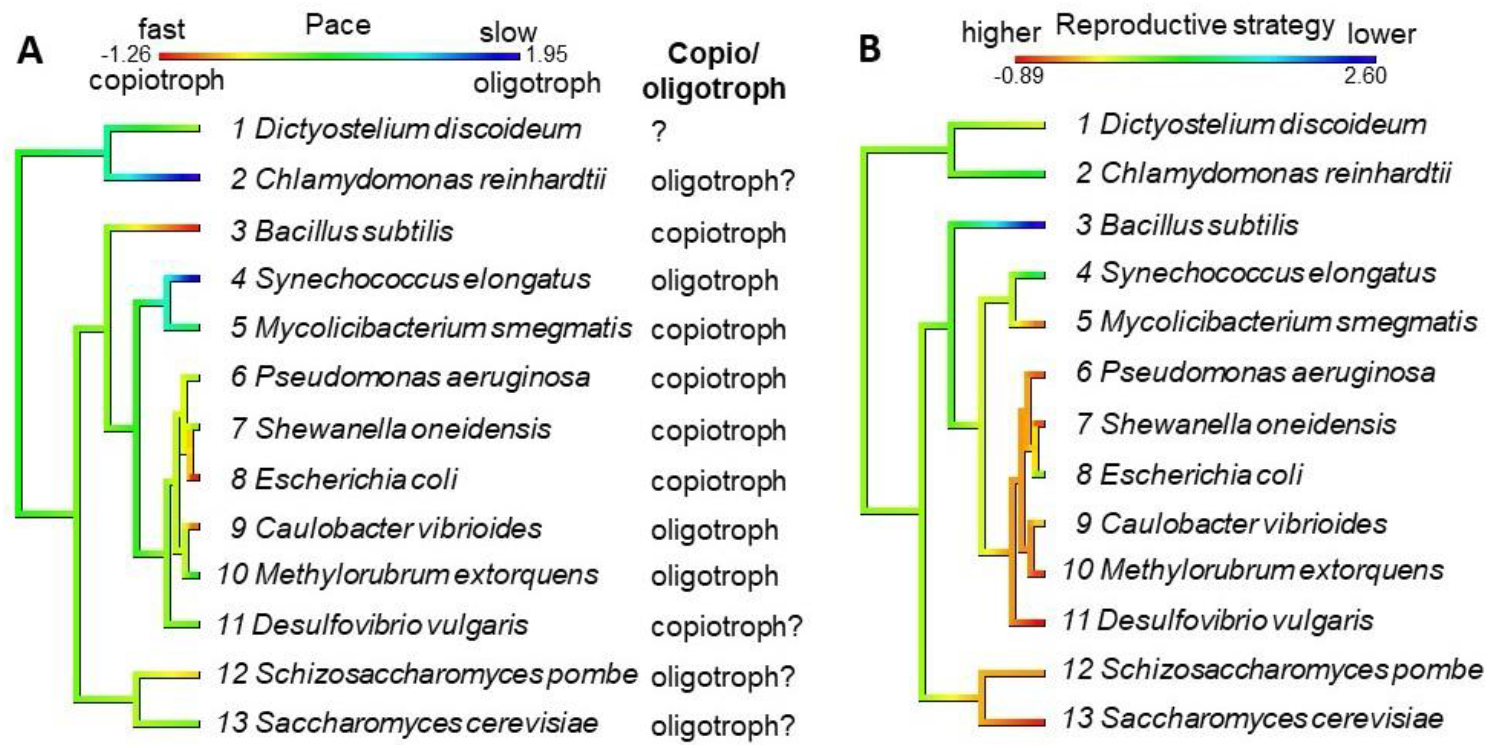
Traitgrams indicating the relationship between a species’ phylogenetic position (as in Fig. 1) and (**A**) its place along the pace of life continuum from fast (negative values in red) to slow (positive values in blue), and (**B**) its place along the reproductive strategy axis from high success (negative values in red) to low success (positive values in blue). Values in (A) are the scores of p-PC1 and values in (B) are the scores of p-PC2 (Fig. 2A). Copiotroph and oligotroph classification was according to [58] (species 1), [5] (species 2-10), [59] (species 11), and [60] (species 12-13) (questions marks indicate suspected classifications).

## Discussion

Microorganisms are typically classified as copiotroph or oligotroph but it has been proven difficult to generalise their life history strategies to broad lineages [5]. Here, we assessed if the fast-slow reproductive strategy framework, which has so far only been applied to animals and higher plants [7-9,35], can be applied to microorganisms. We also tested if inferences about life history strategies can be made from patterns in taxonomic composition of microorganisms. Like animals and higher plants, we found that microorganismal life history variation was structured along a fast-slow pace of life continuum and an axis of reproductive strategy [6,9]. But only at the extreme ends of the pace of life continuum was there a match between copiotrophs at the fast end and oligotrophs at the slow end of the continuum. Our explorative work therefore suggests that the fast-slow reproductive strategy framework can provide a strong predictive framework on the behaviour and activity of microorganism that can to some extent be used alongside the copiotroph-oligotroph dichotomy.

Like previous life history strategy analyses on animals and higher plants, we found that our findings of microorganismal life history strategies were unaffected by phylogeny. Possibly, plasticity in life history traits (variation of trait values within a species that occurs in response to variability in the environment) is obscuring the phylogenetic signal. For example, microorganisms can switch from fermentation to respiration in response to changes in glucose availability, resulting in great reprogramming of gene expression (e.g., [36]). Also, cell growth in e.g., *S. cerevisiae* and *B. subtilis* depends on food quality [23,37], creating plasticity in growth. When variation among individuals in a species is sufficiently large to approach variation between species, any correlation between species traits and their evolutionary relatedness can be masked. Alternatively, or additionally, the lack of a phylogenetic signal may be due to high life history trait lability (e.g., [38]), or due to weak links between genotype and phenotype evolution mapping as life history traits are typically determined by multiple genes that interact to produce phenotypes in a non-additive or non-linear fashion (e.g [39]). If indeed phylogenetic relationships play a minimal role in determining life history adaptations of microorganisms, environmental filtering stemming from extrinsic factors should be considered as the main process of the evolution of microorganismal life history strategies [1].

The evolution of life history strategies is also shaped by the trade-offs that individual make between investment into survival, growth and reproduction [1]. Firstly, the pace of life continuum in animals and higher plants is underpinned by a trade-off between fast-growing, highly reproductive species and population turnover [7,9]. We also found that the microorganismal pace of life continuum was underpinned by a trade-off between highly reproductive species and population turnover, but the highly reproductive species were associated with low values of progressive growth (γ), and, thus, fast microorganisms were slower, and not faster growers. Perhaps, this is because individual microorganisms grow by enlarging individual cells, whereas macroorganisms, among others, grow by increasing total cell mass [40]. Secondly, a trade-off between current and future reproduction underlies the reproductive strategy axis [41,42]. Highly semelparous species with a single reproductive event in their lifespan and high mortality are ranked at one extreme, and iteroparous species with high spread in reproduction and low mortality on the other [10]. We did find, like animals and higher plants, that the reproductive strategy axis captured variation in net reproductive rate (R_0_), but we modelled a semelparous life history strategy for all the microorganisms in our study (as an individual cell dies to produce an offspring and parent cell) [23,43,63]. However, the plant *C. reinhardtii*, the fungi *S. pombe* and *S. cerevisiae*, and the protist *D. discoideum* can reproduce sexually [45-47]. Also, when reproducing asexually, *S. cerevisiae* divides via budding, producing a smaller offspring cell and a remaining, larger parent cell. The plant *C. reinhardtii*, in turn, undergoes a multiple-fission cycle, potentially producing up to eight daughter cells [48]. Future endeavours could explore how more complex growth and reproduction patterns affect life history strategy structuring.

In conclusion, we identified a pace of life and reproductive strategy axis in microorganisms, albeit for a small number of species. Next steps should at least (i) target more species, including from the field,

(ii) systematically control abiotic factors like temperature, and (iii) explore a variety of growth and reproduction patterns. We hope that our first exploration will ignite such research. It could help us understand the diversity of microorganismal phenotypes [49-51], and increase our understanding of how they impact on higher level processes like host phenotype expression [52,53], and human physiology and metabolism [12,13].

## Author statements

## Author contributions

J.R. contributed to data collection, formal analysis, visualisation, and writing of the first draft. I.M.S. contributed to conceptualisation, supervision and writing of final draft.

## Acknowledgements

The authors thank Tom van Dooren for constructive comments on an earlier draft.

